# Knockdown of the translocon protein EXP2, reduces growth and protein export in malaria parasites

**DOI:** 10.1101/420034

**Authors:** Sarah C. Charnaud, Rasika Kumarasingha, Hayley E. Bullen, Brendan S. Crabb, Paul R. Gilson

## Abstract

Malaria parasites remodel their host erythrocytes to gain nutrients and avoid the immune system. Host erythrocytes are modified by hundreds of effectors proteins exported from the parasites into the host cell. Protein export is mediated by the PTEX translocon comprising five core components of which EXP2 is considered to form the putative pore that spans the vacuole membrane enveloping the parasite within its erythrocyte. To explore the function and importance of EXP2 for parasite survival in the asexual blood stage of *Plasmodium falciparum* we inducibly knocked down the expression of EXP2. Reduction in EXP2 expression strongly reduced parasite growth proportional to the degree of protein knockdown and tended to stall development about half way through the asexual cell cycle. Once the knockdown inducer was removed and EXP2 expression restored, parasite growth recovered dependent upon the length and degree of knockdown. To establish EXP2 function and hence the basis for growth reduction, the trafficking of an exported protein was monitored following EXP2 knockdown. This resulted in severe attenuation of protein export and is consistent with EXP2, and PTEX in general, being the conduit for export of proteins into the host compartment.

## Introduction

Almost half the world’s population is at risk of malaria, the disease caused by infection with *Plasmodium spp*. parasites. In 2016 there was an estimated 216 million cases reported, resulting in 445,000 deaths, mostly of children under 5 [1]. *Plasmodium* parasites invade erythrocytes and remodel them to obtain supplementary nutrition from the blood plasma and to evade the immune system. Symptomatic malaria disease is caused by intracellular blood stage parasites which are enveloped in a parasitophorous vacuole membrane (PVM) within the erythrocyte. Residing on the PVM is an essential protein translocon called PTEX (Plasmodium translocon for exported proteins) [2]. PTEX appears to be responsible for exporting hundreds of parasite effector proteins across the PVM into the host erythrocyte where they perform their functions [3, 4].

PTEX consists of five core components, HSP101, PTEX150, EXP2, TRX2 and PTEX88 [2]. Of the core PTEX components only two have homology to other known proteins outside the *Plasmodium* genus. The first is HSP101, a AAA+ heat shock protein chaperone which is predicted to form a hexameric structure to unfold proteins for export [2, 5]. The second is TRX2, a thioredoxin-like protein possibly involved in regulating PTEX or reducing the disulfide bonds in cargo proteins prior to export. TRX2 is not essential for blood stage growth in the murine malaria species *Plasmodium berghei* since its gene can be deleted, however its loss reduces export efficiency and virulence [4, 6, 7]. PTEX150 bears no obvious homology with other proteins, and deletion mutants indicate it is essential and probably responsible for maintaining the structural integrity of PTEX [8]. PTEX88 is a predicted β-propeller protein and appears to be involved in parasite sequestration as knockout or knockdown of PTEX88 in *P. berghei* resulted in reduced sequestration and virulence [9, 10], and an inducible knockdown in *P. falciparum* resulted in reduced binding to the endothelial receptor CD36 [10].

The final core PTEX protein is EXP2 and it is thought to form the PVM spanning pore through which exported proteins are extruded into the erythrocyte compartment [2]. Although EXP2 lacks predicted transmembrane spanning domains typical of membrane pores, it is the most membrane associated protein of the PTEX complex and recombinant EXP2 was observed to form pores in lipid bilayers [2, 11, 12]. Very recently a partial structure of the PTEX complex was solved based on cryo-EM images derived from purified parasites complexes [13]. This indicated seven EXP2 protomers form a funnel-shaped channel in the PVM projecting into the parasitophorous vacuole (PV) lumen. A HSP101 hexamer is anchored via its C-terminus to the EXP2 funnel with seven PTEX150 protomers nestled between the EXP2 protomers helping to form a protein-translocating channel through the center of the structure. Cycles of HSP101 allosteric movements powered by ATP hydrolysis appear to push unfolded proteins through the channel into the erythrocyte via a ratchet mechanism [13].

In addition to a full size PTEX complex of >1200 kDa, we have shown EXP2 forms homo-oligomers of approximately 600 and 700 kDa in size by blue native polyacrylamide gel electrophoresis raising the question of what the smaller forms could be doing [12]. Could EXP2 be a non-selective, high density pore present in a predominantly open state on the PVM that allows low molecular weight compounds of <1400 Daltons across the PVM [14, 15]? Data has shown that EXP2 could complement deletion of GRA17, a predicted pore-like protein in the related Apicomplexan parasite, *Toxoplamsa gondii* [16]. The fact that GRA17 is thought to form a PVM nutrient pore raises the possibility that EXP2 may act in both a large protein translocon complex and in a smaller nutrient import pore [2, 17]. Strongly supporting EXP2’s role as both a protein translocon and as a nutrient channel are recently published patch-clamp experiments with EXP2 knockdown parasites indicating there is reduced conductance of the PVM upon EXP2 knockdown [18].

Export of effector proteins can be inhibited by disrupting the PTEX complex by inducibly knocking down expression of HSP101, PTEX150 and EXP2 [4, 18] or by disrupting the oligomerisation of HSP101 [3]. Loss of PTEX function leads to rapid parasite death within a cell cycle. EXP2 is probably essential for blood stage proliferation since conditional deletion of its gene in *P. berghei* liver stages severely impaired development and patency to blood stage infection [19]. Prior to export, proteins require unfolding and inhibiting this appears to impede PTEX by preventing the export of other essential effector proteins [5, 20, 21].

To explore the role and essentiality of EXP2 in the asexual blood stages of *P. falciparum* we have inducibly knocked down EXP2 expression. We show that EXP2 knockdown is detrimental for parasite growth and the extent of growth arrest is correlated with the degree of knockdown. Upon restoration of EXP2 expression parasite growth can partially recover dependent on the length and strength of knockdown. Upon EXP2 knockdown growth arrest is probably due to the inhibition of export of essential effector proteins since the export of a protein marker is also curtailed.

## Materials and Methods

### Plasmid Construction

From 100 ng of 3D7 *P. falciparum* genomic DNA, the 3’ end of the EXP2 coding region was amplified with forward primer EXP2_1F: AGGAGATCTGGTCACGTATGTGGTGGGTA and reverse primer EXP2_2R A’: CTTATACTGCAGCTTCTTTATTTTCATCTTTTTTTTCATTTTTAAATAAATCTCCACTGGCA. After digestion with *Bgl*II and *Pst*I the *exp2* DNA fragment was ligated into pPTEX150_HAglmS after the *ptex150* flank was removed by digestion with the same enzymes to produce pEXP2-HAglmS.

### Parasite culture and transfection

*P. falciparum* was cultured in human RBCs (Australian Red Cross Blood Bank, blood-group O+) at 4% haematocrit (HCT) in AlbuMaxII media (RPMI-HEPES, 0.5% AlbuMaxII [GIBCO], 0.2% NaHCO_3_, 0.37 mM hypoxanthine) at 37°C as described previously [22]. 100 µg of pEXP2-HAglmS was electroporated into erythrocytes in which *P. falciparum* strain 3D7 was subsequently cultured [23]. Transfected parasites were selected with 2.5 nM WR99210 and were cycled on and off the drug to select for integration into the *exp2* locus. Once western blot validation with anti-HA IgG indicated the *exp2* locus had been appended with the HAglmS construct, clonal parasite lines were isolated by limiting dilution. Successful integration was validated by performing diagnostic PCRs with genomic DNA isolated from each clonal parasite line using the primers: A TGCAGAAACAACTTTGCCACA; A’ TGGCATCTTCTTCTTCAACGGT; B’ TCCTTACGGCTGTGATCTGC.

### EXP2 Knockdown and Western blotting

To knockdown protein expression in EXP2-HAglmS parasites 1M glucosamine (Sigma) was added at the desired concentration to 5 mL of 1% synchronous ring stage parasites in 4% hematocrit. Parasites were grown for 1 cell cycle (∼48 h) and then harvested by centrifugation. The cell pellet was treated with 0.09% saponin in PBS with Complete Protease Cocktail Inhibitors (Roche) to remove the haemoglobin. Cell proteins were solubilised in reducing SDS sample buffer and fractionated on 4-12% acrylamide Bis-Tris gels (Invitrogen) and transferred via iBlot to nitrocellulose membranes (Invitrogen). Membranes were blocked in 1% casein and probed with mouse EXP2 monoclonal, rabbit anti-ERC IgG and chicken anti-HA IgY (Abcam) as per [4]. The membranes were then probed with 700 or 800 nm goat anti-rabbit or goat anti-mouse secondary IgGs (Rockland) and goat anti-chicken IgG-HRP conjugate (Abcam). Probed blots were imaged with a Li-Cor Odyssey InfraRed system and densitometry was performed with Li-Cor Image Studio software.

### EXP2-HA knockdown growth and recovery assays

For parasite growth and recovery assays 5 mL ring stage parasites at 4% hematocrit and 3% parasitemia were treated with the desired GlcN concentration for 1 day. Thin blood smears were then made and 3 x 100 µL aliquots of culture removed for LDH assays. The remaining culture was treated with 0.09% saponin with Complete Protease Cocktail Inhibitors (Roche) to remove the haemoglobin. Parasites to be sampled 2 and 3 days after GlcN treatment were set up as above but diluted 1/5 in 4% hematocrit and those to be sampled after 5 days were diluted 1/25. At days 2 and 3 post GlcN treatment samples were removed for analysis as above. For parasites that were to be grown for 5 days they were feed on days 2 and 3 fed with fresh media and washed to remove GlcN or fresh GlcN was added back. After completion of the time course lactate dehydrogenase (LDH) assays that correlate with parasitaemia were performed as described previously [24] and absorbance measured at 650 nm on a Multiskan GO (Thermo Scientific). The absorbance values were multiplied by culture dilution factors to derive cumulative growth levels at OD 650 nm. Thin blood smears were stained with Giemsa to visualize parasite cell cycle stages. Western blots of EXP2 and ERC levels were performed as described above.

### Microscopy

IFAs were performed as described previously [4]. Parasites were synchronized with sorbitol and heparin to within a 4 h window and plated into 6 well plates at 0.5% parasitaemia and 2% HCT, and GlcN was added at ring stage. One cell cycle later the parasites were collected at approximately 8 hpi and were attached to poly-L-lysine coated slides. The cells were fixed in 4% paraformaldehyde and 0.0075% glutaraldehyde [25], followed by quenching and permeabilisation with 125 mM glycine pH7, 0.05% TX-100 in PBS. Primary antibodies were added at 1:100 in 3% BSA for 4 h. Secondary antibodies (AlexaFluor 488 and 594) were added at 1:2000 for 2 h. Imaging was performed on a Zeiss Axio Observer ZV1 and analysed with FIJI software. Foci of SBP1 were counted automatically as described previously [4]. For counting the ratio of parasite stages, GlcN was added as above and at the appropriate times post GlcN addition, dried blood smears were fixed in 100% methanol for 5 min and stained in 10% Giemsa in water for 30 min. The parasite stages were then counted and photographed.

## Results

### Inducible reduction of EXP2 expression reduces parasite growth

EXP2 is most likely essential for the survival of *P. falciparum* in *in vitro* culture since attempts to genetically delete its gene were not successful in the related rodent parasite *P. berghei* [2, 19]. Given it was not possible to infer EXP2 function in *exp2* null parasites, an alternative approach was adopted where an inducible knockdown system based on the *glmS* ribozyme was employed [4, 26]. In this system the 3’ UTR of *exp2* was substituted with a heterologous 3’ UTR containing *glmS* (Fig. 1A). A haemagglutinin (HA) epitope tag was also appended to the 3’ end of the *exp2* coding sequence for antibody-based detection of the protein. In this knockdown system the *glmS* ribozyme self cleaves the mRNA upon addition of glucosamine (GlcN) to the parasite culture, leading to removal of the poly-A tail, destabilization of the mRNA and subsequent reduction in protein production. The EXP2-HAglmS construct was introduced into the *exp2* locus of the wildtype 3D7 parasite line by homologous recombination (Fig. 1A) [27]. Clonal populations of the EXP2-HAglmS parasites were selected by limiting dilution and integration was confirmed by PCR in six clones (Fig. 1B). Protein knockdown and growth was assessed in clones A12, B12 and G1 and no difference was observed so all subsequent assays were performed with clone B12.

**Figure 1.**
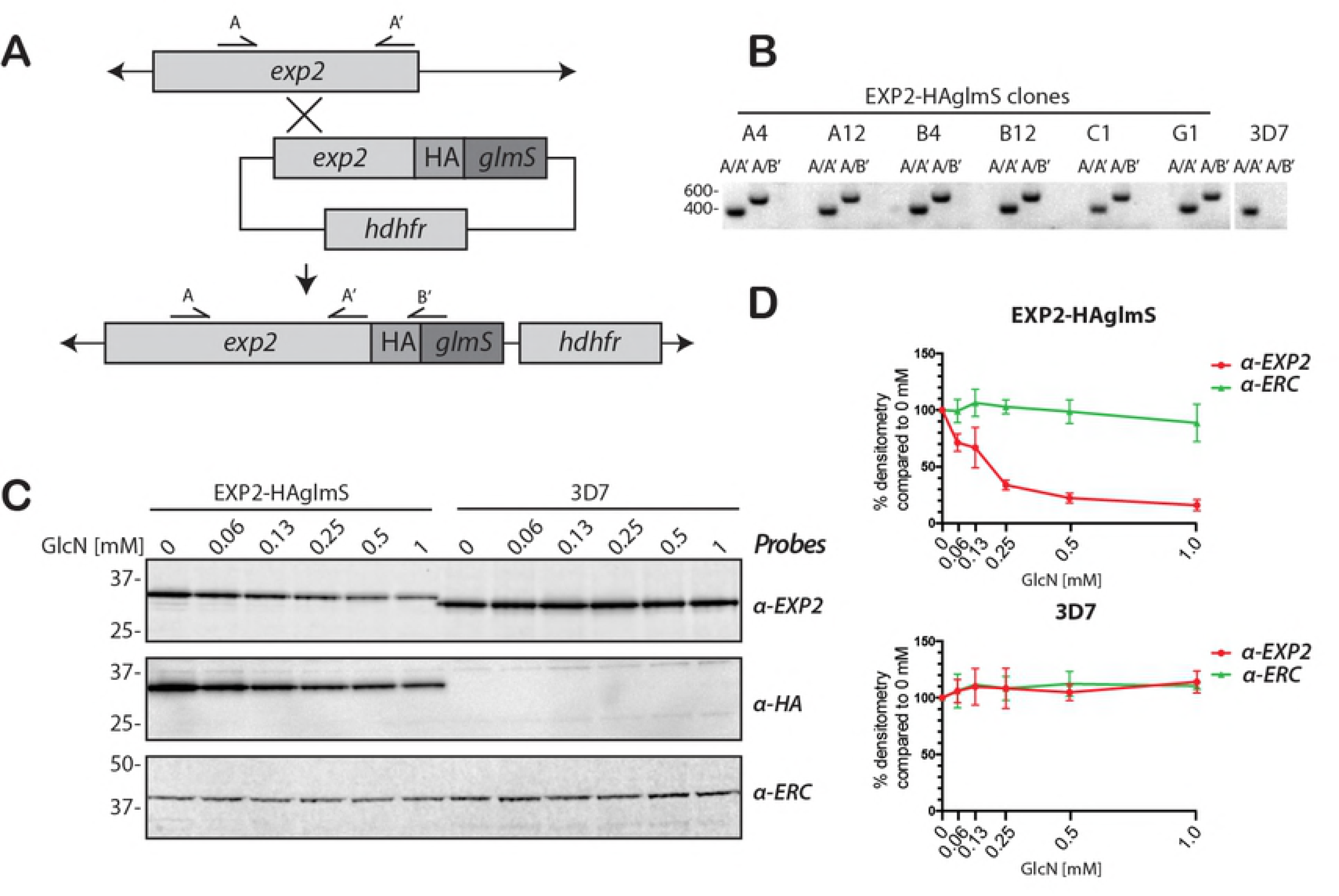
Modification of the exp2 locus with a *glmS* ribozyme permits knockdown of EXP2 expression levels. **(A)** Schematic showing HAglmS tag integrated into exp2 locus. Addition of glucosamine (GlcN) leads to self-cleavage and degradation of mRNA. **(B)** Using the indicated primers and genomic DNA template from wildtype 3D7 and six transfected clonal populations, correct 3’ recombination leading to the HA-*glmS* tagging of *exp2* locus is confirmed. **(C)** GlcN dependent knockdown of EXP2 in EXP2-HAglmS ring stage parasites but not in 3D7 parasites after one cell cycle of GlcN treatment. Size shift in EXP2 indicates HA tagging and is confirmed with a HA-specific antibody. **(D)** Densitometry of EXP2, and ERC loading control from three western blot replicates of GlcN treated EXP2-HAglmS and 3D7 ring stage parasites. Protein densitometry is shown relative to 0 mM GlcN treated parasites normalised to 100%.

To determine the effect of GlcN addition on EXP2 expression, GlcN was added to ring stage parasites at approximately 12-16 hours post invasion (hpi), before expression of EXP2 peaks at 20-25 hpi. Parasite samples were harvested at mid ring stage of the next cycle approximately 48 h later. Subsequent western blots were first probed with a mouse monoclonal for EXP2 that labeled a single band in wildtype 3D7 parasites of about 30.5 kDa predicted for the mature EXP2 protein (Fig. 1C) [2]. A slightly larger band was labeled in EXP2-HAglmS parasites consistent with the addition of the small HA epitope tag (Fig. 1C). This was confirmed with anti-HA chicken IgY that labeled a single similarly sized band in the EXP2-HAglmS parasites but not 3D7 (Fig. 1C). A cross-reactive band of 40 kDa was observed in both parasites that was not therefore specific for the HA epitope (Fig. S1). The blots were also probed for ERC (endoplasmic reticulum-located, calcium-binding protein) to ensure equal protein loading [28]. Examination of the EXP2 and HA blots indicated that GlcN treatment led to a concentration dependent reduction in EXP2 expression in EXP2-HAglmS parasites but not in wildtype 3D7 parasites (Fig. 1D). Quantification of three western blots showed a mean reduction of 84.0% (SD, 4.8%) in EXP2 expression with 1 mM GlcN, similar to that observed for PTEX150-HAglmS [4].

It was expected that a reduction in EXP2 levels would have major effects on the growth and protein export of the parasites on the basis that reducing the function of the other two main PTEX components, PTEX150 and HSP101, had a significant effect [3, 4]. To investigate this, GlcN was added to ring stage EXP2-HAglmS parasites (cycle 0, Fig. 2A). Giemsa stained images of the parasites indicated that maturation into trophozoites in cycle 0 (day 1), and rings of the following cycle 1, (day 2) was not adversely affected (Fig. 2A,B). However by the trophozoite stage of cycle 1 (day 3), GlcN concentrations of 1 and 2 mM had caused parasite growth to arrest. Treatment with 0.5 mM GlcN appeared to have an intermediate effect (Fig. 2A,B). As previous counts of GlcN treated 3D7 parasites indicated no deleterious effects after one cell cycle they were not included here [4].

**Figure 2.**
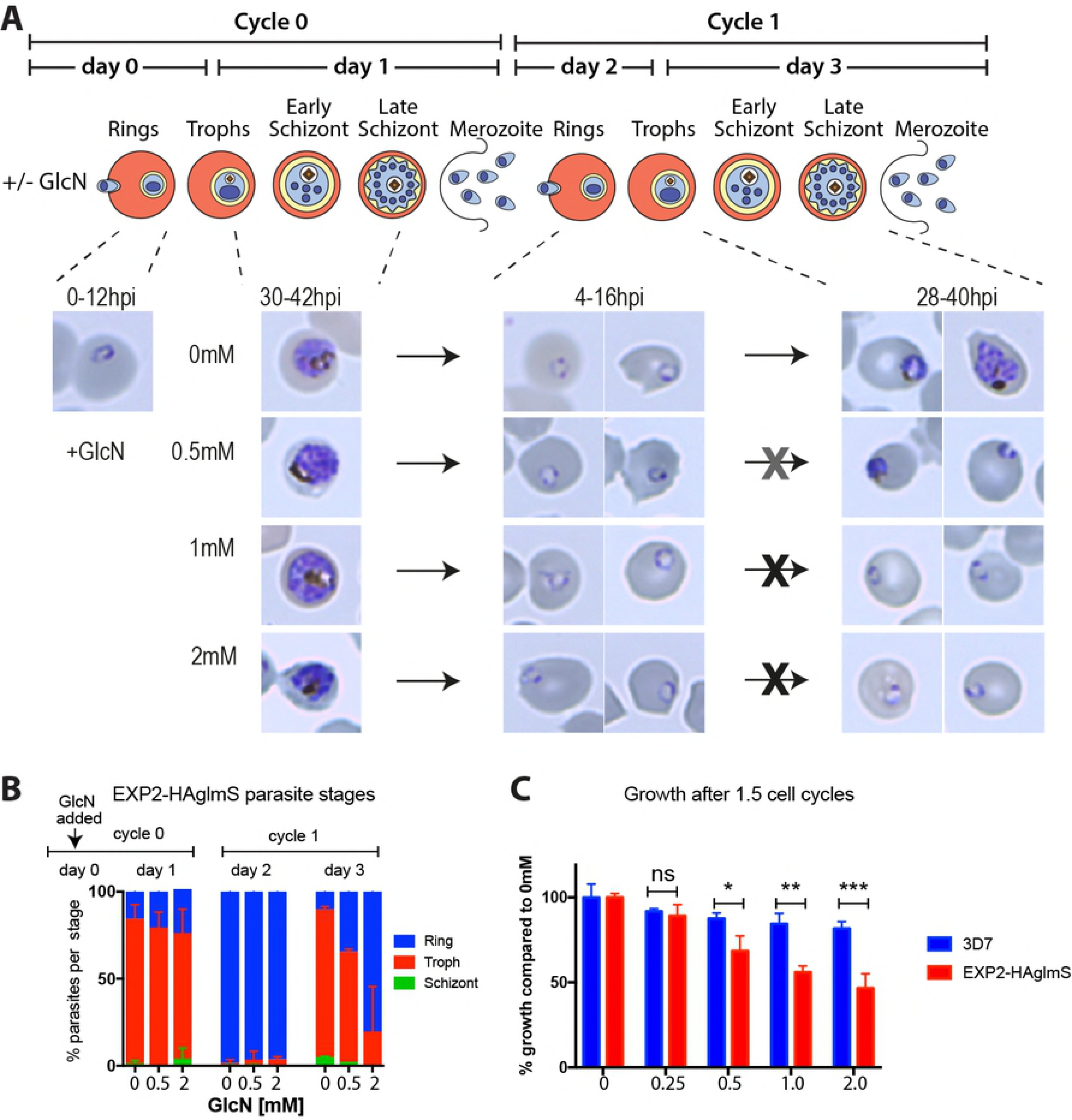
EXP2 knockdown results in a reduced capacity to progress past the ring stage. **(A)** Schematic showing glucosamine (GlcN) addition and EXP2 synthesis. After addition of GlcN in cycle 0 rings, Giemsa smears show that in cycle 1 parasites enter ring stage but appear to stall at the early trophozoite stage. **(B)** Percent of each parasite stage following addition of GlcN to EXP2-HAglmS ring stage parasites at day 0, cycle 0. Giemsa smears of parasite stages were quantified every day for three days (bars indicate mean, SD, representative data of n=2). **(C)** Growth assay after 1.5 cycles measured by lactate dehydrogenase activity (LDH) with increasing GlcN concentrations (n=4). GlcN was added to rings in cycle 0 and LDH activity was measured at trophozoites in cycle 1. Mean, SD are indicated. *p<0.01, **p<0.001, ***p>0.0001.

Next, parasite proliferation using lactate dehydrogenase (LDH) activity as a marker of biomass was measured and indicated that GlcN induced knockdown of EXP2 proportionally reduced proliferation after 1.5 cycles (Fig. 2C). Importantly, GlcN did not significantly reduce the growth of wildtype 3D7 parasites compared to EXP2-HAglmS under any GlcN concentration, indicating it was the knockdown of EXP2 expression that was restricting parasite growth (Fig. 2C).

### Restoration of EXP2-HA expression and parasite growth recovery depends on the degree and length of knockdown

To further explore the relationship between EXP2 levels and parasite growth, 0.5 and 2 mM GlcN that had been added to rings in cycle 0 was washed out in cycle 1 at rings (2 day GlcN treatment) or trophozoites (3 day GlcN treatment). The parasites were then further grown until cycle 2 (5 day) and compared with untreated parasites (0 mM GlcN) and with parasites continuously treated with GlcN for 5 days. Parasites samples from days 1, 2, 3 and 5 were retained for analysis and LDH assays indicated almost full growth recovery after 5 days for parasites treated with 0.5 mM GlcN for 2 and 3 days (Fig. 3A,B). For parasites treated with 0.5 mM GlcN for 5 days, LDH growth was intermediate. Densitometry of the EXP2-HA protein over the growth period from parasites treated with 0.5 mM GlcN indicated the protein was knocked down relative to untreated parasites but quickly recovered once GlcN was removed (Fig. 3C,D). The ERC loading control protein was expressed in near normal levels throughout suggesting that near normal levels of parasites were present, however microscopy of the parasites indicated 5 day GlcN treatment slowed development in some cells leading to the culture becoming asynchronous (Figs 3C,D and S2). In a separate replicate of this experiment we included a 3D7 control and this indicated that 0.5 mM GlcN treatment had no deleterious effect on parasite growth after 5 days (Fig. S3). Western blots were not performed since treatment of 3D7 parasites for 1 cycle (2 days) did not reduce EXP2 or ERC expression (Fig. 1C,D).

**Figure 3.**
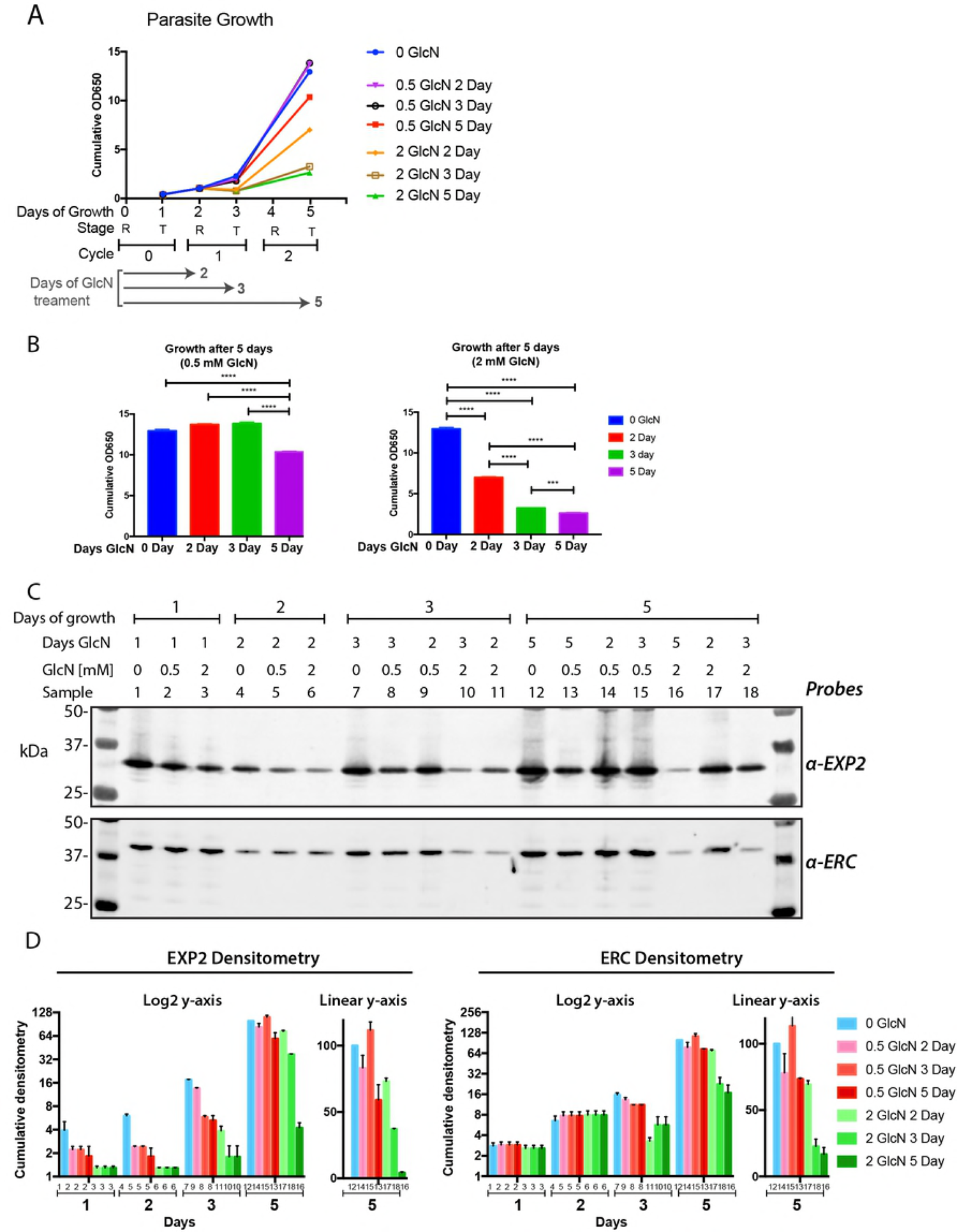
After EXP2-HA is knocked down parasite restoration of growth is dependent upon EXP2 expression. **(A)** GlcN was added to 3% ring stage EXP2-HAglmS parasites to 0, 0.5 and 2 mM on day 0, cell cycle 0. To prevent the parasites from over-growing those that were to be harvested at cycle 1 and 2, were diluted 1/5 and 1/25, respectively at the start of the assay. The parasites were grown for 5 days (2 cell cycles) with samples removed for growth and western blot analysis on day 1 (cycle 0), day 2 and day 3 (cycle 1) and day 5 (cycle 2). To assess recovery of growth after EXP2 knock down, some of the parasites were only treated with GlcN for 2 and 3 days after which the GlcN was removed and growth continued until day 5. To quantify growth, parasite lactate dehydrogenase (LDH) activity of 3 technical replicates was measured on each day. The LDH values (OD 650 nm) were multiplied by the parasite dilution factor to produce a cumulative measure of growth. **(B)** Growth at day 5 after continuous GlcN treatment compared to 2 and 3 days GlcN treament and recovery until day 5. Cumulative OD650 values taken from **(A)**, indicate that growth recovery is higher the shorter the GlcN treatment period and the lower the concentration used. Tukey’s multiple comparison,*** p<0.001, ****p<0.0001. **(C,D)** Western blot analysis and cumulative densitometry of parasite samples taken each day as indicated in **(A)**, shows that EXP2 knockdown is proportional to the GlcN concentration used and the degree of growth inhibition. The loading control ERC, is proportional to parasite biomass indicated by LDH activity and tends to trail behind the degree of EXP2 knockdown. One representative western blot of two is shown. Graph columns indicate mean ± range of n=2. The numbers directly below each column correspond to the western blot sample lanes from which the densitometry was measured. The densitometry of each sample was multiplied by the degree to which the samples were diluted at the start of the assay to produce a cumulative densitometry.

For parasites treated with 2 mM GlcN, parasite growth was severely curtailed after 5 days treatment (Fig. 3A,B). Relative to untreated parasites, growth recovered to intermediate levels if GlcN treatment only lasted 2 days, but barely at all after 3 days treatment. Western blot analysis indicated there was strong knockdown of EXP2-HA after 1 day of 2 mM GlcN treatment but this partly recovered if the GlcN was removed after 2 days (Fig. 3C,D). ERC loading indicated strong reduction in parasite biomass by day 3 which partly recovered after 5 days if the GlcN had been washed out after 2 days. Microscopy of the 2 mM GlcN treated parasites indicated their progression through the cell cycle was greatly delayed and treatment for 3 days probably lead to some parasite death due to the appearance of small dark pyknotic forms (Fig. S2). Control experiments with 3D7 parasites treated with 2 mM GlcN for 5 days slightly reduced growth relative to untreated parasites but not nearly as severely as for EXP2-HAglmS parasites (Fig. S3). It therefore appears that parasites can tolerate a reduction of EXP2 expression for 2 days as long as the level of knockdown was not too great (0.5 mM compared with 2 mM GlcN).

### Knockdown of EXP2-HA reduced export of the skeleton binding protein 1 marker protein

In light of EXP2′s putative PTEX pore function we expected that the growth arrest observed upon reduction of EXP2 levels would be due to the parasite′s inability to export critical proteins into the erythrocyte. To determine if protein export in the EXP2 knockdown parasites was curtailed we performed microscopy on skeleton binding protein 1 (SBP1), an early exported protein, in ring stage parasites before growth arrested. Once exported, SBP1 accumulates at Maurer’s clefts which are parasite-formed vesicular structures that play a role in *Pf*EMP1 export [29, 30]. Immunofluorescence microscopy of 3D7 parasites indicated there were several bright, punctate SBP-labeled structures in the erythrocyte cytoplasm of each cell (Fig. 4A). These were automatically counted using FIJI software and it was found that GlcN treatment did not reduce the average number of SBP1 puncta although their numbers did vary from one treatment to the next possibly due to the numbers of puncta in focus during image acquisition.

**Figure 4.**
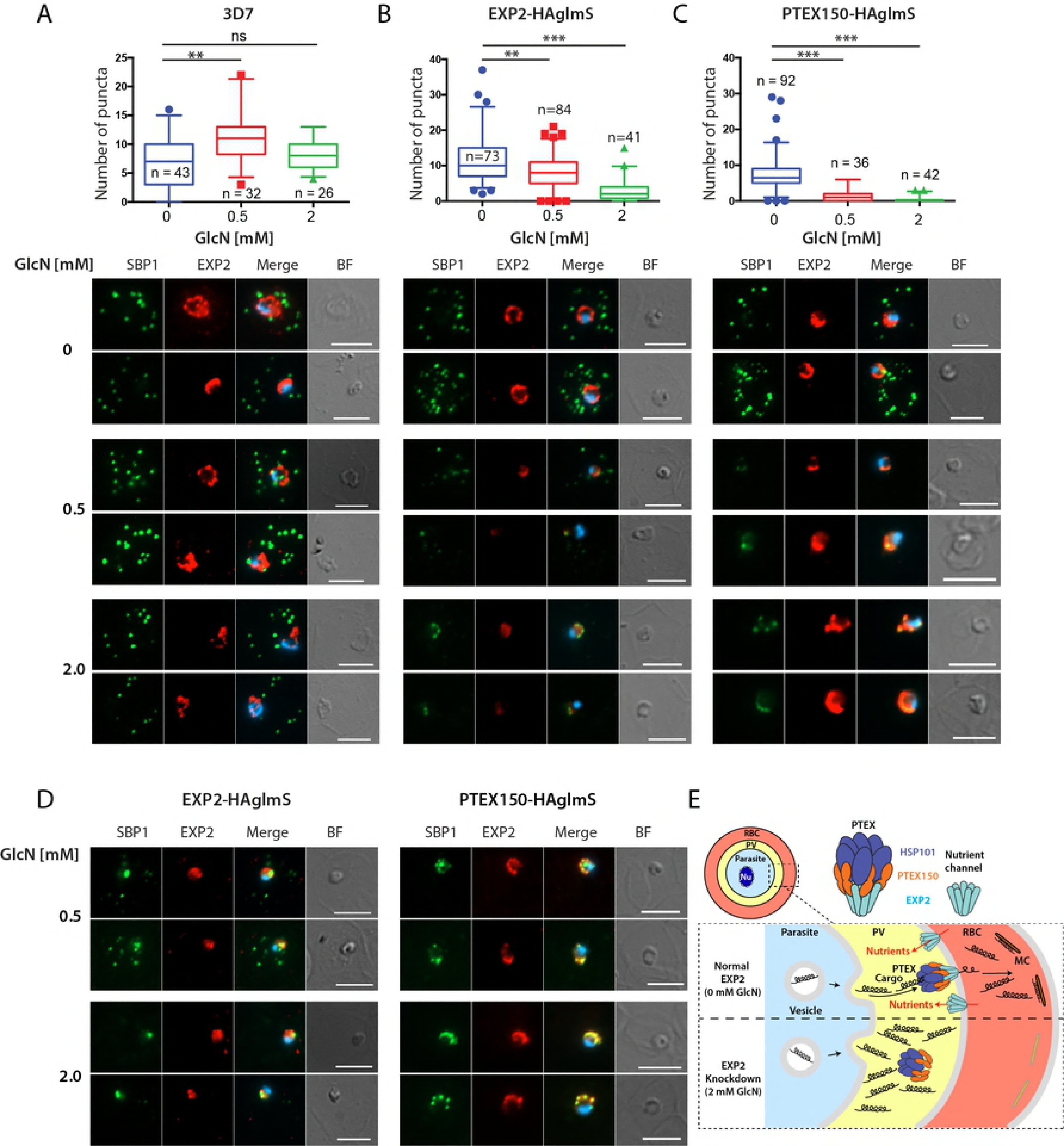
Knockdown of PTEX proteins EXP2 and PTEX150 reduces protein export into host erythrocytes. Export of Maurer’s cleft SBP1 protein after one cycle of GlcN treatment at 8 hpi rings by widefield microscopy in **(A)** 3D7, **(B)** EXP2-HAglmS and **(C)** PTEX150-HAglmS parasites. **(Top)** Quantitation of the number of SBP puncta in the parasites from n = number of individual cells counted. Kruskall-Wallis test used *p<0.01, **p<0.005, *** p < 0.0001. **(Bottom)** Representative immunofluorescence images showing reduced export of SBP structures after treatment of parasites with 0, 0.5 and 2 mM GlcN. Parasites were probed with rabbit anti-SBP-1 serum (green), EXP2 mouse monoclonal IgG (red) and merged with DAPI DNA stain (blue). BF, brightfield. The brightness and contrast for SBP and EXP2 were equally adjusted for each image based on 3D7, 0 mM GlcN. **(D)** Images of GlcN-treated EXP2-HAglmS and PTEX150-HAglmS parasites in which the SBP and EXP2 signal has been enhanced. All size bars = 5 µm. **(E)** Diagram showing how knockdown of EXP2 blocks protein export. PTEX structure derived from Ho et al. (2018)[13]. RBC, red blood cell; PV, parasitophorous vacuole; Nu, nucleus; MC, Maurer’s cleft.

The exposure times required to produce satisfactory images of the untreated 3D7 parasites were used for all subsequent imaging and likewise all brightness and contrast settings were identically adjusted for the SBP1 and EXP2 fluorescent channels. Imaging of EXP2-HAglmS parasites indicated that EXP2 levels were markedly reduced following GlcN treatment (Fig. 4B). Furthermore the number and brightness of SBP1 puncta was also reduced particularly in the 2 mM GlcN treatment. The observed puncta tended to be clustered around the periphery of the parasite and were not exported (Fig. 4B). To determine if SBP1 might still be exported but not visible due to reduced expression the brightness and contrast were increased for SBP1 and EXP2 and this revealed strong trapping of SBP1 at the parasite periphery overlapping with residual EXP2 in 2 mM GlcN (Fig. 4D). In 0.5 mM GlcN there was also SBP1 trapping but some export was observed.

Although we have previously reported GlcN mediated export trapping in PTEX150-HAglmS parasites we repeated this here for comparison with EXP2-HAglmS [4]. GlcN treatment of PTEX150-HAglmS parasites did not reduce EXP2 expression as anticipated but did decrease the intensity of SBP1 puncta and caused them to overlap with EXP2 around the parasite periphery (Fig. 4C). Enhancement of the SBP1 signal revealed that export of virtually all this protein was blocked. Casual observation of the both parasite lines suggested that export was more efficiently blocked in PTEX150-HAglmS compared to EXP2-HAglmS particularly in 0.5 mM GlcN. This was supported by the SBP1 puncta counts that indicated a stronger reduction in PTEX150-HAglmS (Figs 4B,C).

## Discussion

To validate an essential role for EXP2 in the PTEX protein export complex we attempted to knockdown EXP2 and examine subsequent phenotypes since gene knockout attempts have not produced a population of viable parasites that could be phenotypically studied [2, 19]. Our knock down strategy also appended a HA tag and *glmS* ribozyme onto EXP2 and reduced its expression by up to 84% following treatment with 1 mM GlcN. This appeared to arrest parasite growth in late rings one cycle after adding GlcN to degrade the protein’s mRNA. The EXP2-HA knockdown parasites export SBP1 less efficiently than parasites with normal levels of EXP2 suggesting growth arrest could be due to the failure to export functionally crucial proteins other than SBP1, which itself is not essential [29, 30]. During preparation of this work another study was published in which EXP2 was knocked down by 90% using a TetR-DOZI-aptamer system [18]. Both this study and ours similarly confirm EXP2 knockdown greatly reduced parasite growth as well as protein export. Although our study does not shed light on the pore forming role of EXP2, two recent studies confirm that it serves both as a protein pore for the PTEX complex and as a potential nutrient pore for transfer of low molecular weight solutes across the PVM [13, 18].

That the *glmS* mediated knockdown of EXP2-HA was responsible for growth reduction is provided by the fact that the HAglmS tag only appeared to append to EXP2 and no other genes since no other HA bands apart from EXP2 were present in western blots. The degree of knockdown of EXP2-HA protein was critically dependent on the concentration of GlcN added to the parasite culture and GlcN had no significant effect on 3D7 parasites. Furthermore, when GlcN was removed EXP2 expression and parasite growth were partially restored. Specifically, when 0.5 mM GlcN was washed out after 2 and 3 days treatment, EXP2 protein levels and parasite proliferation almost fully recovered by day 5. At a higher dose of 2 mM GlcN for 2 days, only partial recovery occurred by day 5. GlcN mediated knockdown of EXP2 also greatly reduced export of SBP1 consistent with EXP2 being part of the PTEX complex [29, 30]. Confirmation that EXP2 is part of the PTEX complex was also provided by a recent separate study, where immunoprecipitation of EXP2-HA pulled down all four other proteins of the PTEX complex as well as a cargo reporter protein [21].

Our initial data suggested that growth of EXP2-HAglmS parasites was arrested by 0.5 mM GlcN treatment after 2 - 3 days, however closer investigation revealed that growth was more likely just slowed given that the parasites recovered well once GlcN was removed and EXP2 expression resumed. These parasites had about a half to a third the normal amount of EXP2-HA after 2 – 3 days treatment and could probably still export enough proteins and obtain sufficient nutrients to survive and hence recovered following GlcN removal. The cultures also appeared to become less synchronous suggesting that the ∼48 h cell cycle was extended in some cells. Whether slowed growth also resulted in fewer merozoites being formed was not examined, but nutrient restriction via knockdown of the new permeability pathway protein RhopH2 has been noted to reduce merozoite numbers per schizont [31], therefore this may also be a factor underpinning growth delay in GlcN-treated EXP2HAglmS parasites.

Parasites seem to be only able to tolerate the strong knockdown of EXP2 caused by 2 mM GlcN treatment for a few days since after this growth recovery was poor and dead parasites began to be observed after 3 days of 2mM GlcN treatment. After 2 days of 2 mM GlcN treatment EXP2 levels had fallen to 20% of normal levels and this enabled many parasites to survive since they partially recovered by day 5 if the GlcN was washed out after 2 days. After 3 days 2 mM GlcN treatment when levels of EXP2 had fallen to ∼10%, recovery was poor probably because most parasites had died by then. The ability of ring stage arrested parasites to partially recover growth as long as 2 days after arrest has been previously detected for another PTEX protein HSP101, and is consistent with our EXP2 observations here [3].

Both our study and Garten et al (2018) [18] showed that the PEXEL negative exported protein (PNEP) SBP1 export is reduced when EXP2 is knocked down [18]. We are confident that the trafficking of other exported proteins, in particular PEXEL proteins, would also be similarly inhibited since we have seen this previously following the knockdown of PTEX150 [4]. Knockdown of HSP101 expression in *P. berghei* also reduced export of several different PEXEL proteins and in *P. falciparum* knockdown of HSP101 function reduced the export of several PEXEL and PNEP proteins [3, 4]. A knockout study of ∼50 exported proteins has indicated that most PEXEL proteins (∼75%) are probably non-essential including most of those listed above [30, 32-34]. This raises the question of what essential functions are carried out by exported proteins whose export blockage results in parasite growth arrest? Although the intra-erythrocytic locations of many exported proteins are known along with their effects on erythrocyte rigidity, knob formation, *Pf*EMP1 display and the trafficking of other exported proteins, the precise functional role for most exported proteins is not understood, with a few exceptions [35]. A recent attempt to carry out saturation mutagenesis of the *P. falciparum* genome with piggyback transposons has helped determine which genes might be unmutable and essential and are therefore a priority for future study [36]. Although the essentiality of PTEX might depend on the critical functions of a small number of essential exported proteins it could also be equally dependent on minor functions of dozens of non-essential proteins that collectively have a huge effect on parasite viability when their trafficking is blocked.

In conclusion, we have shown that EXP2 is crucial for survival of *P. falciparum* blood stages probably because it is required for the efficient export of functionally important proteins into the erythrocyte compartment. Knockdown of EXP2 might be additionally slowing growth by restricting the availability of nutrients to the parasite and the removal of wastes.

## Acknowledgments

We thank the Australian Red Cross Blood Bank for the provision of human blood, Jacobus Pharmaceuticals for providing WR99210 and Monash Micro Imaging. We are grateful to Leann Tilley and Brian Cooke for providing ERC and SBP1 antibodies and Eva Pesce and Thorey Jonsdottir for technical assistance. The authors gratefully acknowledge funding from the Victorian Operational Infrastructure Support Program received by the Burnet Institute and for grants from the National Health and Medical Research Council of Australia (1068287, 1128198 and 1092789).

## Author Contributions

S.C.C., P.R.G and B.S.C designed, managed and funded the project; S.C.C., P.R.G. and R.K. produced experimental data and S.C.C, P.R.G. and H.E.B. wrote manuscript.

## Supporting information captions

**Figure S1. Original full size western blot images use to construct Fig.1C are shown along with the antibodies and fluorescent channels used to detect target proteins.** Note that that the same blot was probed and detected in different channels. The regions of the blots selected for construction of Fig. 1C are indicated with a red dashed line.

**Figure S2. Knockdown of *glmS* tagged *ptex* genes reduces protein expression and inhibits parasite growth.** Giemsa stained thin blood smears showing slowed growth and parasite death caused by knockdown of EXP2-HAglmS. Parasite smears are from experiment shown in Fig. 3.

**Figure S3. Glucosamine treatment greatly reduces the growth of EXP2-HAglmS parasites relative to 3D7 control parasites. (A)** GlcN was added to 3% ring stage parasites to 0, 0.5 and 2 mM on day 0, cell cycle 0. To prevent the parasites from over-growing those that were to be harvested at cycle 1 and 2, were diluted 1/5 and 1/25, respectively. The parasites were grown for 5 days or 2 cell cycles with samples removed for growth and western blot analysis on day 1, cycle 0; day 2 and day 3, cycle 1; and day 5, cycle 2. To assess recovery of growth after EXP2 knock down, some of the parasites were only treated with GlcN for 2 and 3 days after which the GlcN was removed and growth continued until day 5. To quantify growth, parasite lactate dehydrogenase (LDH) activity of 3 technical replicates was measured at each time point. The LDH values (OD 650 nm) were multiplied by the parasite dilution factor to produce a cumulative measure of growth. **(B)** Growth at day 5 after continuous GlcN treatment compared to 2 and 3 days GlcN treatment and recovery until day 5. Cumulative OD650 values taken from **(A)**, indicate that growth recovery is higher the shorter the GlcN treatment period and lower the concentration used. Tukey’s multiple comparison,*** p<0.001, ****p<0.0001.

